# Phase variation of a signal transduction system controls *Clostridioides difficile* colony morphology, motility, and virulence

**DOI:** 10.1101/690230

**Authors:** Elizabeth M. Garrett, Ognjen Sekulovic, Daniela Wetzel, Joshua B. Jones, Adrianne N. Edwards, Germán Vargas-Cuebas, Shonna M. McBride, Rita Tamayo

## Abstract

Recent work has revealed that *Clostridioides difficile*, a major cause of nosocomial diarrheal disease, exhibits phenotypic heterogeneity within a clonal population as a result of phase variation. Many *C. difficile* strains representing multiple ribotypes develop two colony morphotypes, termed rough and smooth, but the biological implications of this phenomenon have not been explored. Here, we examine the molecular basis and physiological relevance of the distinct colony morphotypes produced by this bacterium. We show that *C. difficile* reversibly differentiates into rough and smooth colony morphologies, and that bacteria derived from the isolates display opposing surface and swimming motility behaviors. We identified an atypical phase-variable signal transduction system consisting of a histidine kinase and two response regulators, named herein CmrRST, which mediates the switch in colony morphology and motility behaviors. The CmrRST-regulated surface motility is independent of Type IV pili, suggesting a novel mechanism of surface expansion in *C. difficile*. Microscopic analysis of cell and colony structure indicates that CmrRST promotes the formation of elongated bacteria arranged in bundled chains, which may contribute to bacterial migration. In a hamster model of acute *C. difficile* disease, colony morphology correlates with virulence, and the CmrRST system is required for disease development. Furthermore, we provide evidence that CmrRST phase varies during infection, suggesting that the intestinal environment impacts the proportion of CmrRST-expressing *C. difficile*. Our findings indicate that *C. difficile* employs phase variation of the CmrRST signal transduction system to generate phenotypic heterogeneity during infection, with concomitant effects on bacterial physiology and pathogenesis.

**Significance Statement:** Phenotypic heterogeneity within a genetically clonal population allows many mucosal pathogens to survive within their hosts, balancing the need to produce factors that promote colonization and persistence with the need to avoid the recognition of those factors by the host immune system. Recent work suggests that the human intestinal pathogen *Clostridium difficile* employs phase variation during infection to generate a heterogeneous population differing in swimming motility, toxin production, and more. This study identifies a signal transduction system that broadly impacts *C. difficile* physiology and behavior *in vitro* and in an animal model. Phase variation of this system is therefore poised to modulate the coordinated expression of multiple mechanisms influencing *C. difficile* disease development.

## Introduction

Phenotypic heterogeneity within bacterial populations is a widely established phenomenon that allows the population to survive sudden environmental changes (1–3). Heterogeneity serves as a “bet-hedging” strategy such that a subpopulation can persist and propagate an advantageous phenotype. Phase variation is one mechanism that imparts phenotypic heterogeneity, typically by controlling the ON/OFF production of a surface-exposed factor that directly interfaces with the environment (4, 5). Many human pathogens, including pathogenic *Escherichia coli, Neisseria meningitidis*, and *Streptococcus pneumoniae*, employ phase variation to evade the host immune system and increase host colonization, persistence, and virulence (4, 6–13). Phase variable phenotypes are heritable yet reversible, allowing the surviving subpopulation to regenerate heterogeneity. In phase variation by conservative site-specific recombination, a serine or tyrosine recombinase mediates the inversion of a DNA sequence adjacent to the regulated gene(s) (5). The invertible DNA sequences are flanked by inverted repeats and contain the regulatory information, such as a promoter, that when properly oriented regulates gene expression *in cis*.

*Clostridioides difficile* (formerly *Clostridium difficile*) is a gram-positive, spore-forming, obligate anaerobe and a significant public health burden globally, causing gastrointestinal disease ranging from diarrhea to potentially fatal complications such as pseudomembranous colitis, toxic megacolon, bowel perforation, and sepsis. *C. difficile* pathogenesis is primarily driven by the toxins TcdA and TcdB, which inactivate host Rho-family GTPases resulting in actin cytoskeleton dysregulation, tight junction disruption, and host cell death; consequently, TcdA and TcdB compromise the epithelial barrier and elicit diarrheal symptoms and inflammation (14–16). Many aspects of *C. difficile* physiology and pathogenesis remain poorly understood. For example, some *Clostridioides* species, including *C. difficile*, are capable of forming two distinct colony morphologies; one is smooth and circular, and the other is rough and filamentous (17–20). However, the underlying mechanisms and physiological relevance of this dimorphism are unknown. Many bacterial species develop multiple colony morphologies as a result of the regulated production of surface factors, which can impact diverse physiological traits and behaviors (21–26). In multiple species, the production of surface factors is subject to phase variation, leading to changes in gross colony morphology. In *S. pneumoniae* and *Acinetobacter baumannii*, phase variation of capsule polysaccharides leads to the formation of either opaque or translucent colonies that differ in a multitude of phenotypes including cell morphology, biofilm formation, antibiotic sensitivity, host colonization, and virulence (6, 9, 12, 24, 26, 27). Phase variation of colony morphology is therefore an important adaptive strategy during infection for multiple pathogens.

The biological significance and mechanisms underlying colony morphology development of *C. difficile* have not been reported. *C. difficile* encodes multiple factors that are regulated through phase variation. In *C. difficile*, seven invertible DNA sequences, or “switches”, have been identified (28). Two have been characterized: one controlling phase variation of the cell wall protein CwpV and the other controlling flagellar phase variation (19, 29–31). The phase variable *flgB* flagellar operon encodes the sigma factor SigD, which coordinates flagellar gene expression and positively regulates the *tcdA* and *tcdB* toxin genes (32–34). Consequently, the flagellar switch mediates phase variation of flagella and toxin production, highlighting the potential impact of phase variation on *C. difficile* physiology and virulence.

As in many bacteria, the intracellular signaling molecule c-di-GMP regulates the transition between community-associated and planktonic, motile lifestyles of *C. difficile* (33, 35–38). This regulation occurs in part through direct control of gene expression by c-di-GMP via riboswitches (39–42). For example, a c-di-GMP riboswitch lies in the 5’ leader sequence of the *flgB* operon and causes transcription termination when c-di-GMP is bound, thus inhibiting swimming motility (39, 40, 43). Because flagellum and toxin gene expression is linked through SigD, c-di-GMP also inhibits toxin production (34, 43). Conversely, a c-di-GMP riboswitch upstream of the Type IV pilus (TFP) locus allows c-di-GMP to positively regulate gene expression and promote TFP-dependent behaviors such as autoaggregation, surface motility, biofilm formation, and colonization of host tissues (33, 35–37, 43, 44). Additionally, c-di-GMP regulates the expression and cell wall anchoring of surface proteins and adhesins (41, 45, 46). Therefore, c-di-GMP coordinates the expression of multiple surface factors with implications for pathogenesis.

In this study, we characterized rough and smooth colony isolates of *C. difficile* R20291 and determined that they exhibit distinct motility behaviors *in vitro*. Colony morphology and associated motility phenotypes are controlled by both phase variation and c-di-GMP, and colony morphology is independent of TFP and flagella. We identified the c-di-GMP regulated, phase-variable signal transduction system, consisting of a putative histidine kinase and two DNA-binding response regulators, that modulates colony morphology, surface migration, and swimming motility. Finally, we provide evidence using a hamster model of infection for phase variation and a role for the CmrRST system in virulence. The link between phase variation and c-di-GMP signaling to control CmrRST production suggests a mechanism for switching a global expression program during infection, which appear to have critical implications for disease development.

## Results

### *C. difficile* strains of diverse ribotypes develop two distinct, phase-variable colony morphotypes

The *C. difficile* strain R20291, a ribotype 027 strain associated with epidemic infections, exhibits two distinct colony morphologies: a “smooth” colony that is round and circular, and a “rough” colony that is flatter and has filamentous edges (18–20). In addition to R20291, strains UK1 (ribotype 027), VPI10463 (ribotype 087), 630 (ribotype 012), and ATCC 43598 (ribotype 017) yielded both rough and smooth colonies (Figure 1). Some of these strains (R20291 and UK1) showed both colony morphotypes through routine plating, while others (630) required extended incubation to allow the appearance of bacteria that develop rough colonies. ATCC BAA 1875 (ribotype 078) did not develop smooth colonies under any conditions tested. Therefore, many strains are capable of generating two distinct colony morphologies *in vitro*.

**Figure 1.**
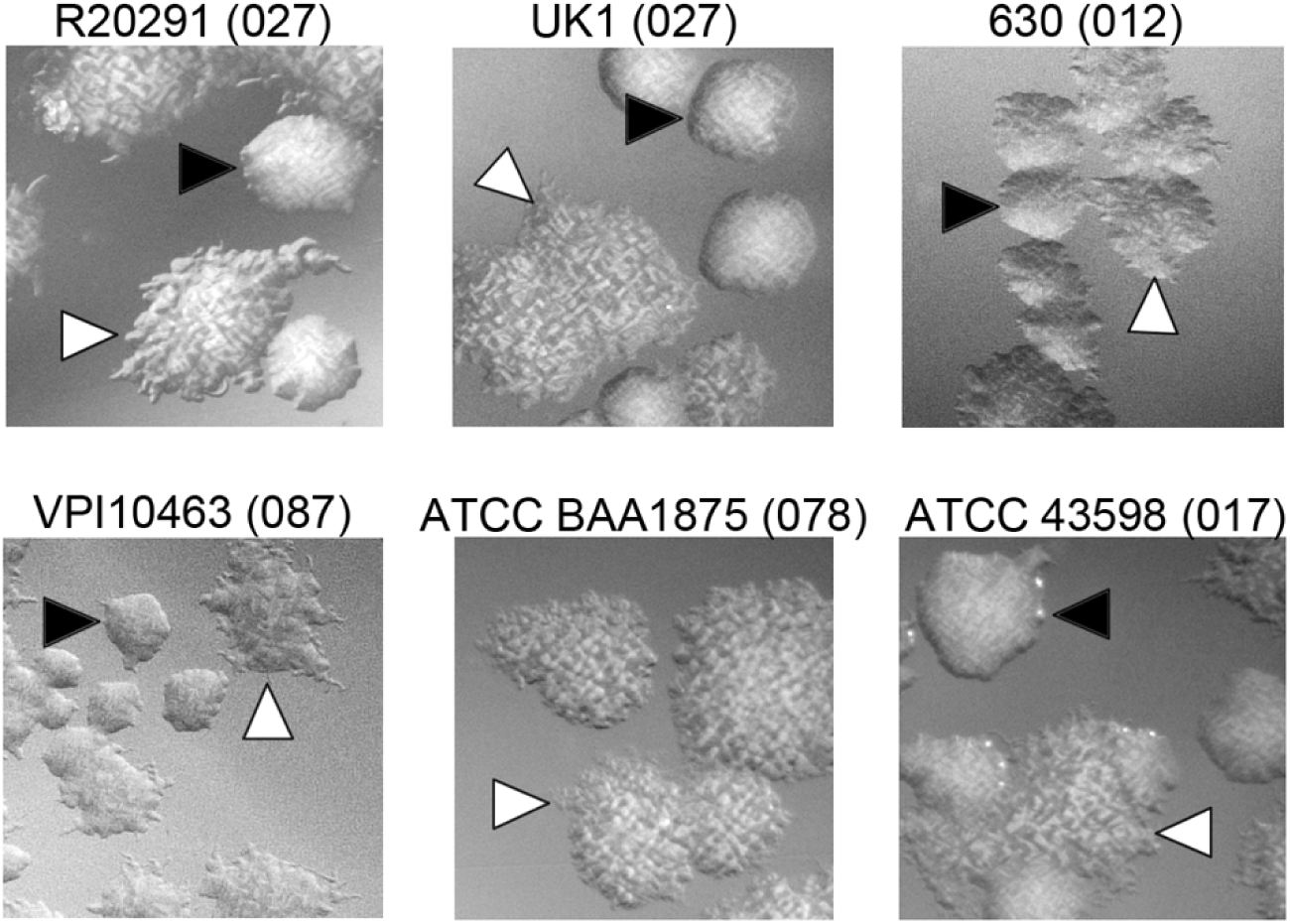
Formation of rough and smooth colonies by multiple *C. difficile* strains. *C. difficile* strains representing five different ribotypes (indicated in parentheses) were grown on BHIS-agar medium for 72 hours to allow differentiation of colony morphology. Growth was recovered and plated on BHIS-agar for visualization of individual colonies. Shown are representative images from two independent experiments. Black triangles indicate “smooth” colonies, and white triangles indicate “rough” colonies. All images were taken at 2X magnification.

We and others previously observed that spotting a culture of *C. difficile* R20291 on an agar surface results in expansion of the colony as dendritic, filamentous growth (18, 33). We discovered that this culturing method allows the segregation of the two colony morphologies. Streaking from the center of the colony yielded a population of predominantly smooth colonies, while streaking from the edges yielded a population of predominantly rough colonies (Figure 2A). To evaluate the stability of the colony phenotypes, we passaged rough and smooth colonies through multiple growth conditions. Streaking a rough or smooth colony again onto agar medium resulted in overall maintenance of the original morphology (Figures 2A,C), indicating that colony morphology is heritable and mostly stable under specific growth conditions. Similarly, the starting morphology was maintained when the isolates were passaged in broth (Figure 2C). Because surface growth promoted the development of the rough morphotype, we speculated that conditions favoring swimming motility might yield smooth colonies. Rough and smooth colonies were inoculated into soft agar to allow for swimming motility over 48 hours (Figure 2B, panel 4). Growth recovered from the edges of the motile spot solely yielded smooth colonies, regardless of the starting morphology, indicating that colony morphology is reversible (Figures 2B,C). These data support that *C. difficile* colony morphology phase varies and reveal *in vitro* growth conditions that select for a specific phase variant: surface growth favors the expansion of bacteria that yield rough colonies, while swimming conditions select for bacteria that yield exclusively smooth colonies.

**Figure 2.**
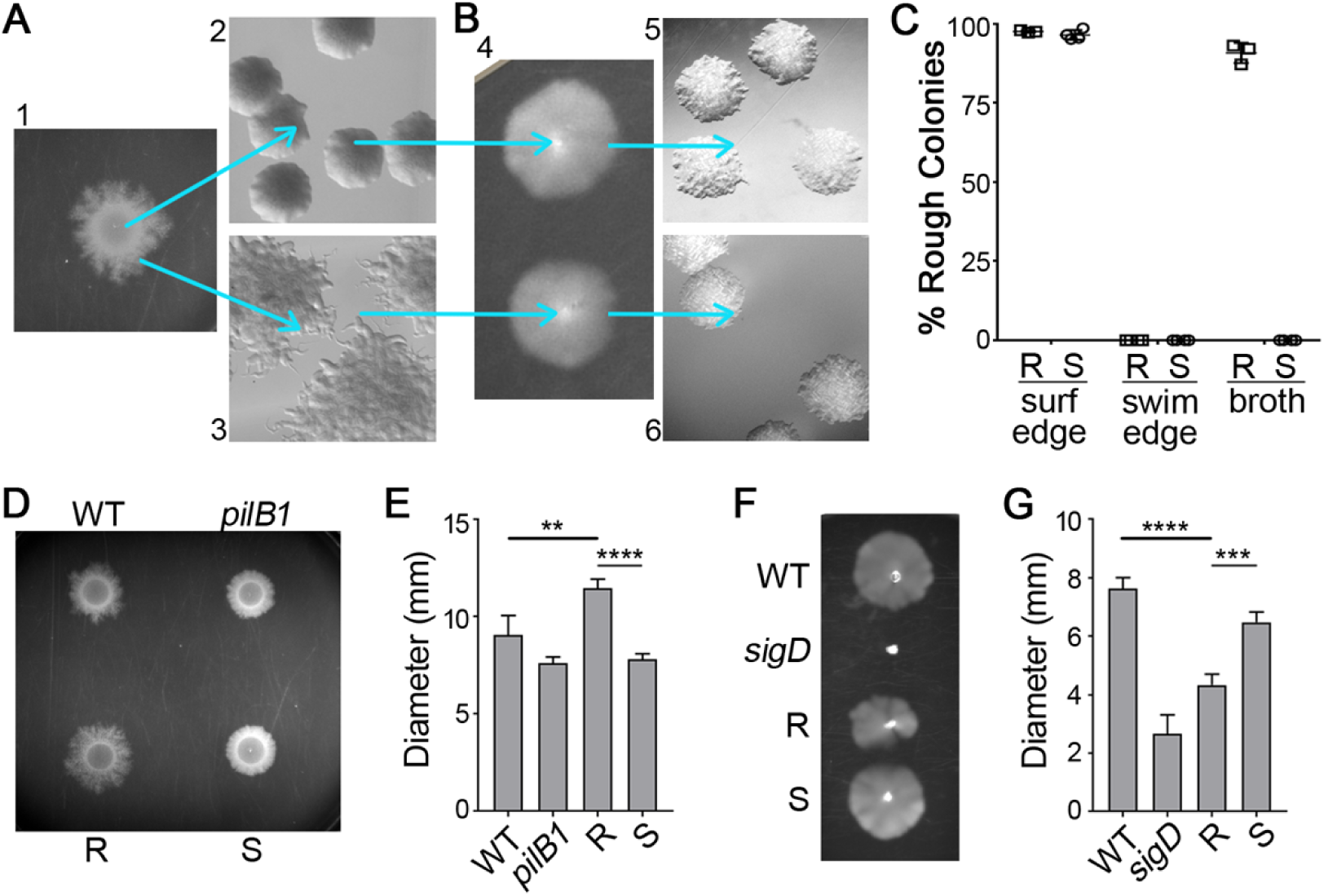
Reversible selection of distinct colony morphotypes with opposing motility phenotypes. (A) *C. difficile* R20291 liquid culture was spotted on BHIS-1.8 % agar and grown for 48 hours to allow spreading growth (panel 1). Bacteria collected from the center of a spot yield mostly smooth colonies (panel 2), while bacteria from the edge yield almost exclusively rough colonies (panel 3). (B) Rough and smooth colony isolates obtained as in A were passaged in 0.5X BHIS-0.3% agar (panel 4), and samples were collected from the edge of motile growth. Both rough and smooth colony isolates only gave rise to smooth colonies (panels 5,6). Shown are representative images of colonies from four independent experiments. All images were taken at 2X magnification. (C) Quantification of colony morphology for rough and smooth colony isolates after passage on a BHIS-agar surface (surf) as in A or in motility medium (swim) as in B, with samples collected from the edges of growth. R, S indicate rough and smooth starting inoculums, respectively. Passage in BHIS broth medium served as a control. Symbols indicate individual values from 3-4 biological replicates, and bars indicate means and standard deviations. (D,E) *C. difficile* R20291, a TFP-null control (*pilB1*), and rough (R) and smooth (S) colony isolates were assayed for surface motility on BHIS-1.8 % agar. (D) A representative of four experiments is shown. (E) Surface motility was quantified by measuring the diameter of growth after 48 hours. (F,G) *C. difficile* R20291, a non-motile control (*sigD*), and rough (R) and smooth (S) colony isolates were assayed for swimming motility in 0.5x BHIS-0.3% agar. (F) A representative of four experiments is shown. (G) Swimming motility was quantified by measuring the diameter of growth after 48 hours. (E,G) **p < 0.01, ***p < 0.001, ****p < 0.0001, one-way ANOVA and Tukey’s post-test.

### Rough and smooth colony isolates exhibit distinct motility behaviors

The observation that surface motility conditions favor bacteria that develop rough colonies suggests that the rough morphotype has an advantage over smooth in this growth condition. Conversely, smooth morphotype bacteria may have an advantage over the rough in swimming motility. To test this, we assessed the motility behaviors of rough and smooth colony isolates. Surface motility of “wild-type” (i.e. undifferentiated by colony morphology) R20291 is characterized by initial (∼ 24 hours) growth restricted to the site of inoculation of an agar surface, then spreading tendrils of growth between 24 and 96 hours (36). Type IV pili contribute to surface motility, so the non-piliated *pilB1* mutant was included as a control (35, 36). In this assay, bacteria isolated from rough colonies exhibited greater surface motility after 48 hours compared to undifferentiated (WT) R20291 (Figures 2D,E). Conversely, bacteria derived from smooth colonies were deficient in surface motility compared to the rough isolates and more similar to the *pilB1* control after 48 hours (Figures 2D,E), though the smooth isolates remained capable of surface motility.

Flagellum-dependent swimming motility was assayed by inoculating bacteria into 0.3% agar and measuring migration through the medium. Undifferentiated R20291 and a non-motile *sigD* mutant served as controls (19). Rough colony isolates showed significantly decreased swimming motility compared to smooth and undifferentiated populations (Figures 2F,G). Bacteria from smooth colonies showed swimming motility comparable to the undifferentiated parent (Figures 2F,G), likely because the parental isolate consisted primarily of bacteria that yield smooth colonies. These results indicate that rough and smooth colony morphotypes have distinct motility behaviors.

### Colony morphology is regulated by a phase variable signal transduction system and c-di-GMP

Because colony morphology development exhibited characteristics suggestive of phase variation, we sought to identify the underlying mechanism. To date, the only known phase variation mechanism in *C. difficile* involves site-specific DNA recombination resulting in a reversible DNA inversion that is known or predicted to impact gene expression *in cis* (28). We postulated that one of the seven known invertible sequences regulates the expression of genes involved in colony morphology development, which would be evident as a correlation between the orientation of the invertible sequence and colony morphology (Figure 3A). To test this idea, we differentiated rough and smooth populations and analyzed the orientation of the “switches” in these isolates using qPCR with orientation-specific primers. For six of the switches, no correlation between morphology and sequence orientation was observed (Figure 3B). However, the orientation of the invertible element Cdi6, which is upstream of the operon CDR20291_3128-3126, showed a strong correlation with colony morphology (28, 30). Each of four independently isolated rough populations contained the sequence predominantly in the “on” orientation previously determined to favor gene expression, while each of the smooth populations contained the sequence predominantly in the inverse “off” orientation (28). CDR20291_3128-3126 encodes a putative phosphorelay system consisting of two predicted response regulators and a predicted histidine kinase (28). Accordingly, we named the operon <u>c</u>olony <u>m</u>orphology regulators RST, where *cmrR* and *cmrT* encode the response regulators and *cmrS* encodes the histidine kinase, and refer to the Cdi6 invertible element as the “*cmr* switch”. The *cmrRST* locus and its upstream regulatory region containing the *cmr* switch are present in all 65 complete *C. difficile* genomes available on NCBI with > 96% identity at the nucleotide level. This high conservation across diverse ribotypes indicates that this regulatory system, as well as the ability to control its production through phase variation, is important for *C. difficile* fitness.

**Figure 3.**
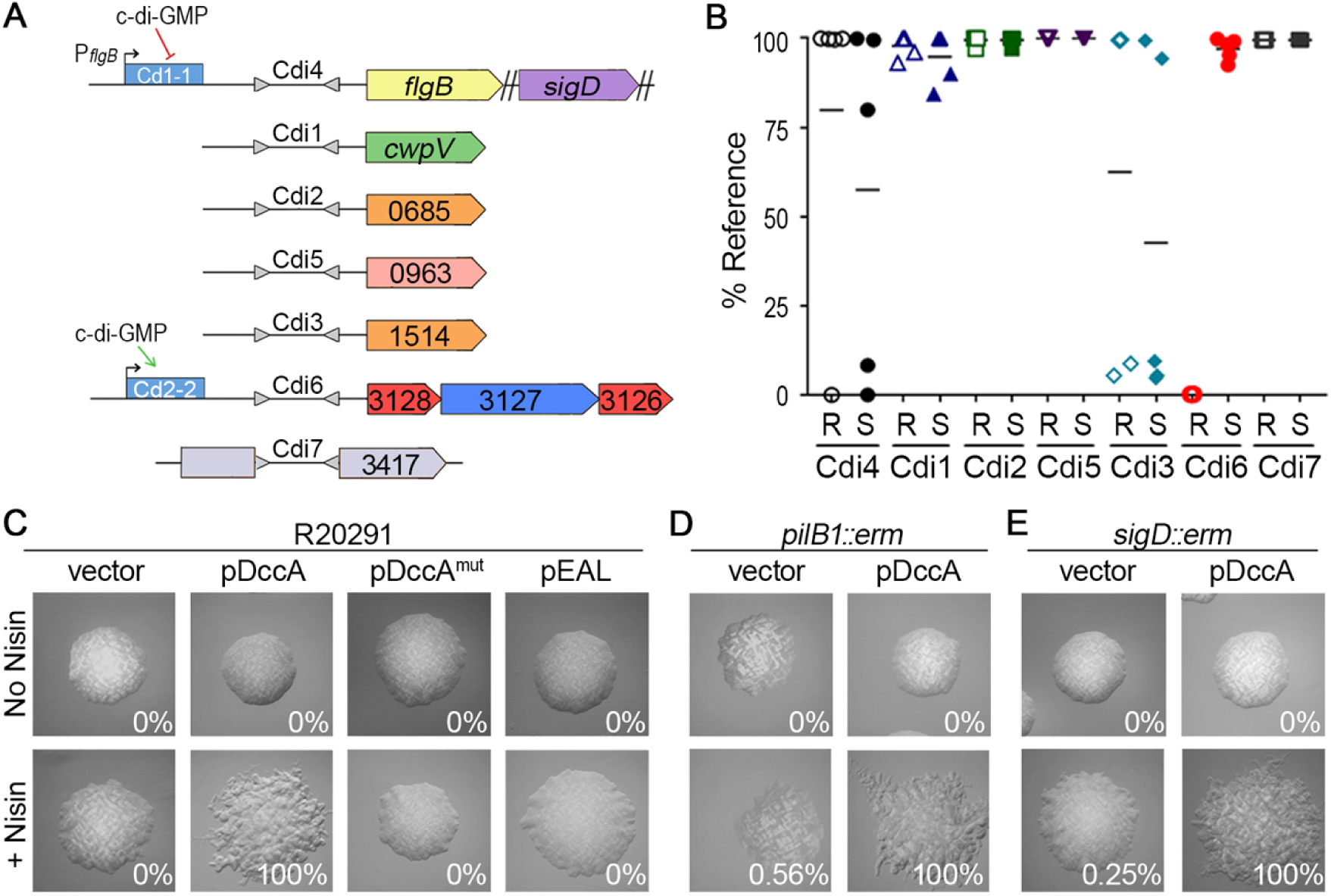
Expression of a c-di-GMP regulated phosphorelay system promotes rough colony formation in a TFP- and flagellum-independent manner. (A) Diagram of seven invertible DNA elements previously identified in *C. difficile* R20291. Gray triangles represent inverted repeats flanking invertible elements, which are named according to previously defined nomenclature [29]. Downstream regulated genes are shown. Blue rectangles denote c-di-GMP riboswitches, and direction of regulation is indicated with arrows. (B) qPCR analysis of the orientations of the seven invertible DNA sequences in rough (R) and smooth (S) colony isolates. Data are expressed as the percentage of the population with the sequence in the orientation present in the reference genome (<u>FN545816</u>). Symbols indicate values from five individual isolates, and horizontal bars indicate the means. (C-E) *C. difficile* R20291, a TFP-null (*pilB1*) mutant, and an aflagellate (*sigD*) mutant containing the indicated plasmids for manipulation of intracellular c-di-GMP were grown on BHIS-agar with or without inducer (2 µg/ml nisin). Shown are images representative of the most common morphology yielded by each strain. Values in each panel indicate the percentage of rough colonies of ≥ 100 total colonies from at least two independent experiments. All images were taken at 2X magnification.

A c-di-GMP binding riboswitch (Cdi-2-2) lies 5’ of *cmrRST* and the invertible sequence and positively regulates expression of the operon (Figure 3A) (39, 41). Additionally, rough and smooth colony morphotypes exhibited inverse swimming and surface motility phenotypes, which is characteristic of c-di-GMP regulation (36, 43). Therefore, we hypothesized that c-di-GMP regulates *C. difficile* colony morphology. Using a previously described strategy, intracellular c-di-GMP was increased or decreased by ectopically expressing genes encoding the diguanylate cyclase DccA or the EAL domain from PdcA, respectively (33, 36). Expression of *dccA* led to the uniform development of rough colonies, while the expression of a catalytically inactive DccA (DccA^mut^) did not, indicating that increased c-di-GMP stimulates rough colony formation (Figure 3C). Reducing c-di-GMP through overproduction of the EAL domain did not impact colony morphology, possibly because basal c-di-GMP levels are already low (33). To determine whether the rough colony effect results from c-di-GMP-mediated inhibition of flagellar motility or activation of TFP-dependent surface motility, *dccA* was expressed in TFP-null *(pilB1::erm*) or aflagellate (*sigD::erm*) backgrounds. The *pilB1* and *sigD* mutants remained capable of forming both smooth and rough colonies in unmodified and increased c-di-GMP conditions (Figures 3D,E). Together, these results indicate dual regulation of the CDR20291_3128-3126 locus by both phase variation and c-di-GMP. Furthermore, c-di-GMP regulates colony morphology through a mechanism independent of TFP and flagella.

### The response regulators CmrR and CmrT regulate colony morphology

Because activation of *cmrRST* expression, whether by increased c-di-GMP and/or inversion of the switch to the “on” orientation, promotes the development of rough colonies, we examined the contributions of the response regulators CmrR and CmrT to this process. The *cmrR* and *cmrT* genes were individually expressed under the control of an ATc-inducible promoter in *C. difficile* R20291 during growth on BHIS-agar. Both CmrR and CmrT stimulated rough colony formation compared to non-induced controls (Figure S2). Expression of *cmrT* resulted in a rough phenotype at lower levels of induction than *cmrR*. Interestingly, some levels of induction of *cmrT* inhibited growth, suggesting that *cmrT* expression is toxic. Growth inhibition by CmrT, but not CmrR, was observed in broth culture as well (Figure S1A, S1B).

Response regulators typically have a conserved aspartic acid residue in the phosphoreceiver domain that, when phosphorylated, leads to activation (47). Mutation of this residue to a glutamic acid often mimics phosphorylation (48, 49). Accordingly, compared to *cmrR* expression, lower levels of induction of *cmrR*^D52E^ expression were required for formation of rough colonies (0.5 ng/ml versus 4 ng/ml ATc, respectively; Figure S2). Instead of a conserved aspartic acid residue, CmrT contains a glutamic acid at the expected phosphorylation site, precluding the need for a phosphomimic substitution, and potentially explaining the lower level of inducer needed to yield rough colonies compared to *cmrR*. Substitution of the phosphorylation site of CmrT with alanine (CmrT-D53A) increased the concentration of inducer needed to obtain rough colonies. However, for the wild type and CmrR-D52A mutant, comparable levels of inducer (4 ng/mL ATc) resulted in rough colonies; this may be due to the need for an activating signal that is absent in these assay conditions. That the inactivating mutations did not abolish CmrR and CmrT function suggests that phosphorylation is not required for the activity of these response regulators if they are expressed at high levels.

Because CmrR and CmrT promoted the rough colony morphotype, we examined the requirement of *cmrR* and *cmrT* in rough colony formation. Individual in-frame deletions of *cmrR* and *cmrT* were generated in *C. difficile* R20291. The mutants and undifferentiated wild-type R20291 parent were grown on BHIS-agar to allow selection of rough colony isolates from the edges (as in Figure 2). While the parent strain and *cmrR* mutant formed both rough and smooth colonies, the *cmrT* mutant did not form rough colonies under the given conditions (Figure 4A). Expression of *cmrT in trans* complemented the *cmrT* mutation, restoring the capacity to form rough colonies (Figure 4B). Importantly, *cmrR* and *cmrT* mutants do not differ in growth rate from WT (Figure S1C). Thus, while CmrR and CmrT both impact colony morphology development when overproduced, only CmrT is required for rough colony formation under the tested conditions.

**Figure 4.**
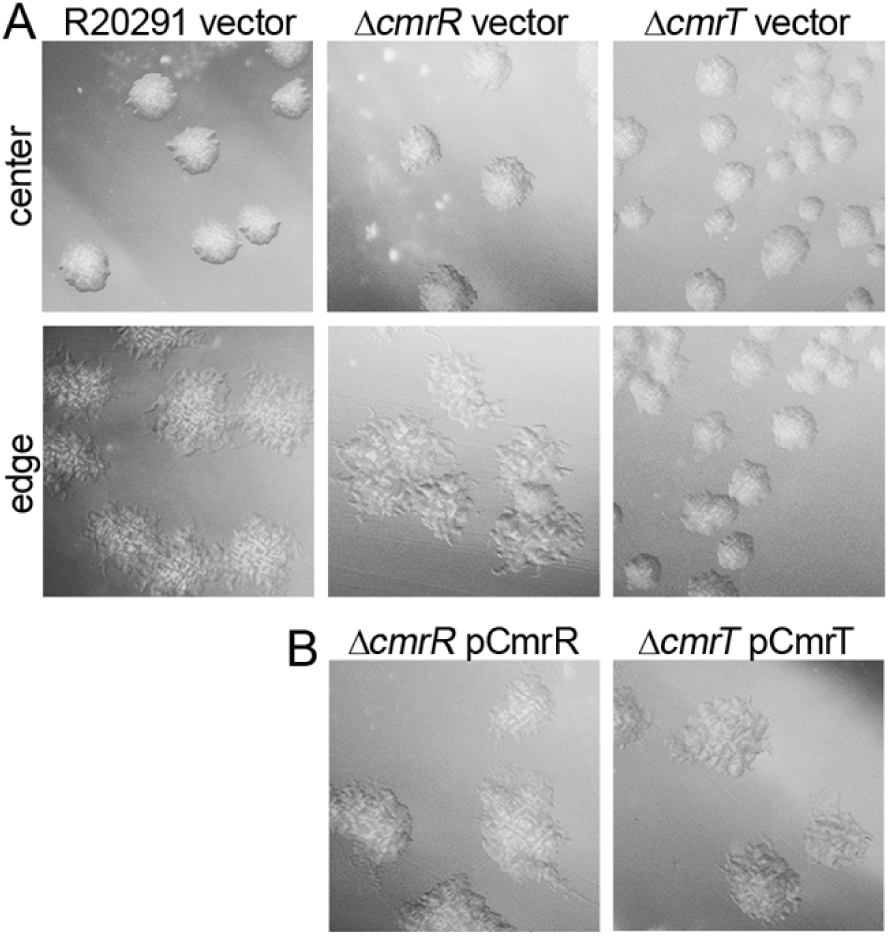
CmrT is required for rough colony formation. (A) Colony morphology of *C. difficile* R20291, Δ*cmrR*, Δ*cmrT* with vector after 24 hours on BHIS-1.8% agar. (B) Complementation of Δ*cmrR* and Δ*cmrT* mutants with expression of the respective genes *in trans* induced with 2 ng/mL ATc, imaged after 24 hours. All images were taken at 2X magnification. Representative images from at least two independent experiments are shown.

### CmrR and CmrT inversely regulate surface and swimming motility

Because bacteria isolated from rough and smooth colonies differed in surface and swimming motility (Figure 2), we assessed the roles of CmrR and CmrT in these processes. Ectopic expression of *cmrR* or *cmrT* in R20291 significantly increased surface motility compared to uninduced and vector controls (Figures 5A, S3A,B). Enhanced surface motility was also observed in a TFP-deficient (*pilB1::erm*) background, indicating that that CmrR and CmrT-mediated surface motility is independent of TFP (Figures 5A, S3A,B). Conversely, *cmrR* and *cmrT* expression inhibited swimming motility of R20291 (Figures 5D, S3C,D). These data are consistent with the increased surface motility and decreased swimming motility exhibited by rough colony isolates (*cmrRST* ON), which are the inverse of the motility phenotypes of the smooth colony isolates (*cmrRST* OFF). Expression of *cmrR* and *cmrT* alleles with inactivating alanine substitutions resembled the wild-type allele, suggesting again that high levels of expression overcome the need for phosphorylation (Figure S3). However, as seen with colony morphology (Figure S2), the *cmrR-*D52E allele increased surface motility and decreased swimming motility at lower levels of induction relative to the wild-type allele, suggesting that the amino acid substitution increased activity (Figure S3). Expression of *cmrR* or *cmrT* did not alter transcript levels of representative flagellum or TFP genes (Figure S4), suggesting that the observed differences in surface and swimming motility occur through post-transcriptional regulation or through an alternative mechanism.

**Figure 5.**
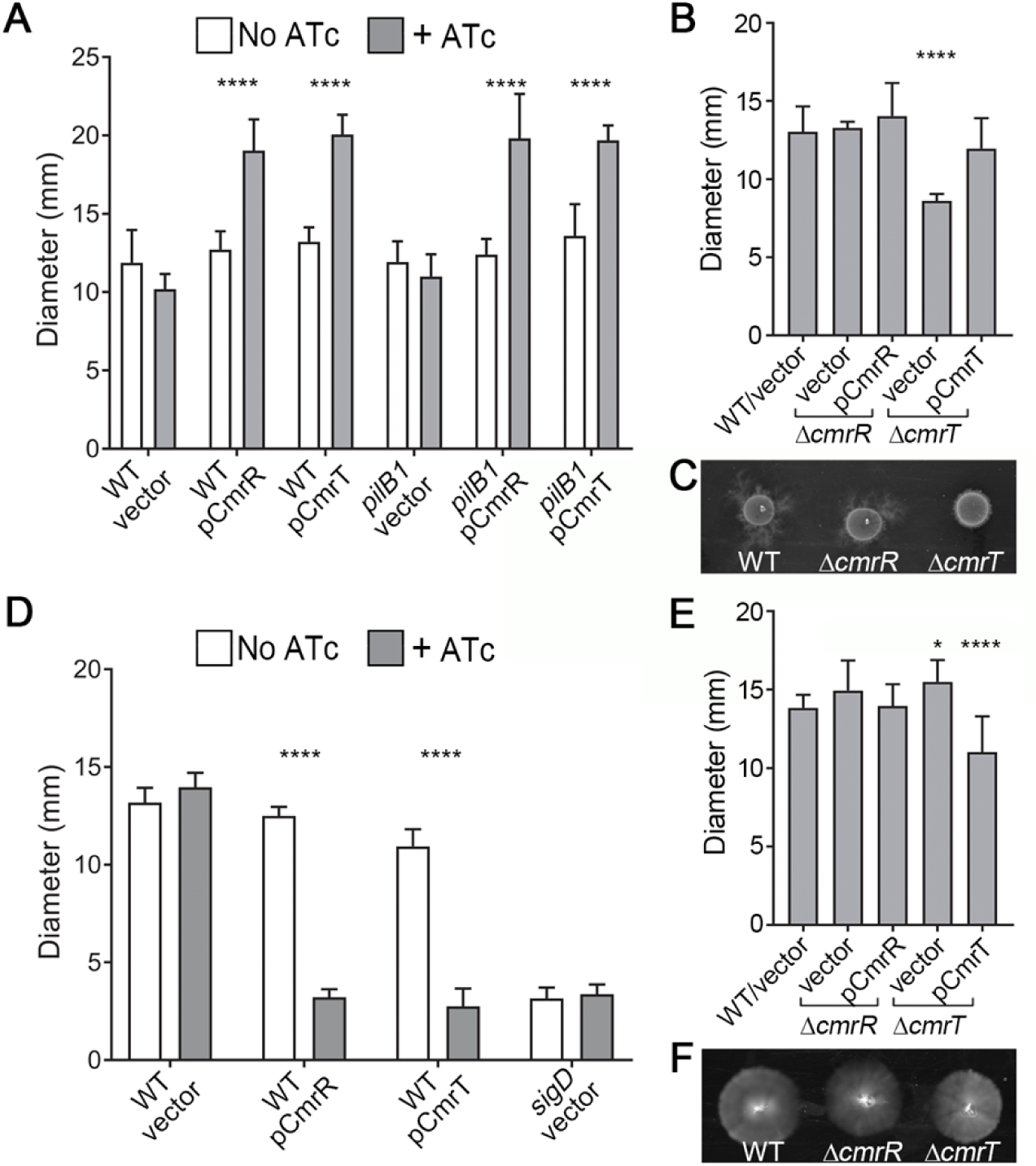
CmrR and CmrT inversely regulate surface and swimming motility. *C. difficile* strains were assayed for surface motility on BHIS-1.8% agar 1% glucose after 72 hours (A-C) and for swimming motility through 0.5X BHIS-0.3% agar after 48 hours (D-F). (A) *C. difficile* R20291 (WT) and a TFP-null (*pilB1*) mutant, each with plasmids for expression of *cmrR*, *cmrT*, or a vector control, assayed for surface motility with or without ATc (10 ng/µL for pCmrR, 2 ng/µL for pCmrT). N ≥ 7 biological replicates combined from four independent experiments. (B) Surface motility of WT, Δ*cmrR*, Δ*cmrT* strains with vector or respective expression plasmids for complementation. N = 16 biological replicates combined from two independent experiments. (C) Representative image of surface motility. (D) *C. difficile* R20291 (WT) and a non-motile *sigD* mutant with pCmrR, pCmrT, or vector as indicated, assayed for swimming motility with or without ATc (10 ng/µL for pCmrR, 2 ng/µL for pCmrT). N ≥ 7 biological replicates combined from four independent experiments. (E) Swimming motility of WT, Δ*cmrR*, Δ*cmrT* strains with vector or complementation plasmids, in the presence of 0.2 ng/µL. A non-motile *sigD* mutant was included as a control. N = 22 biological replicates from three independent experiments. (F) Representative image of swimming motility. (A,B,D,E) Shown are the means and standard deviations of the diameters of motility growth. *p < 0.05, ****p < 0.0001, two-way ANOVA and Tukey’s post-test comparing +/−ATc (A,D) or one-way ANOVA with Dunnett’s post-test comparing values to WT/vector (B,E).

To further define the role of *cmrRST*, the *cmrR* and *cmrT* mutants and relevant controls were evaluated for surface and swimming motility. In both assays, the *cmrR* mutant behaved comparably to the R20291 parent (Figures 5B,C,E,F), indicating that CmrR is dispensable for regulation of surface and swimming motility. However, the *cmrT* mutant exhibited significantly reduced surface motility, a defect that was complemented by expressing *cmrT in trans*. The opposite effect was observed for swimming motility; the *cmrT* mutant showed greater motility compared to the parent strain. CmrT thus appears to be the dominant response regulator of the CmrRST system for control of rough colony formation, surface motility, and swimming motility *in vitro*.

### CmrR and CmrT promote bacterial chaining

Phenotypic analysis of motility and quantification of flagellum and TFP transcripts indicates that CmrRST mediated colony morphology occurs independently of these surface structures. Cell morphology has also been shown to affect gross colony morphology (26, 50, 51), so we investigated whether cell morphology differs between rough and smooth morphotypes. Examination of *C. difficile* colonies by scanning electron microscopy (SEM) revealed the presence of the expected bacillus towards the center, as well as elongated cells particularly along the edge of the colony (Figure 6A). These elongated cells appeared as organized bundles corresponding to the tendrils apparent in the macrocolony, suggesting a role for the elongated bacteria in expansion of rough colonies. Consistent with this, SEM of bacteria derived from smooth and rough colonies revealed dichotomous cellular morphologies. Smooth colony-derived bacteria appeared as standard bacilli (4.089 µm ± 1.207 in length), while rough colony-derived cells were longer (7.428 µm ± 4.130) and sometimes exhibited an extremely elongated shape, resembling a bacterial filament or chain (Figure 6B,C). To determine whether the elongated cells result from filamentation or cell chaining, cells from rough and smooth colonies were Gram stained. While cells from smooth colonies consisted of the expected single or double rods, cells from rough colonies more commonly appeared in bacterial chains of three or more cells (Figure 6E,F). Septa separating cells within chains were clearly visible, differentiating this morphology from filamentation. Differences in processing may explain why cell chains were more common in Gram stained samples than in SEM imaged samples; processing of samples for SEM may have disrupted chains. Since *cmrRST* expression favors rough colony development, we examined the roles of CmrR and CmrT in the chaining phenotype. In contrast to uninduced and vector controls, cells over-expressing *cmrR* or *cmrT* also displayed bacterial chaining (Figure 6G, H), indicating that rough colonies contain bacterial chains whose formation is promoted by either CmrR or CmrT.

**Figure 6.**
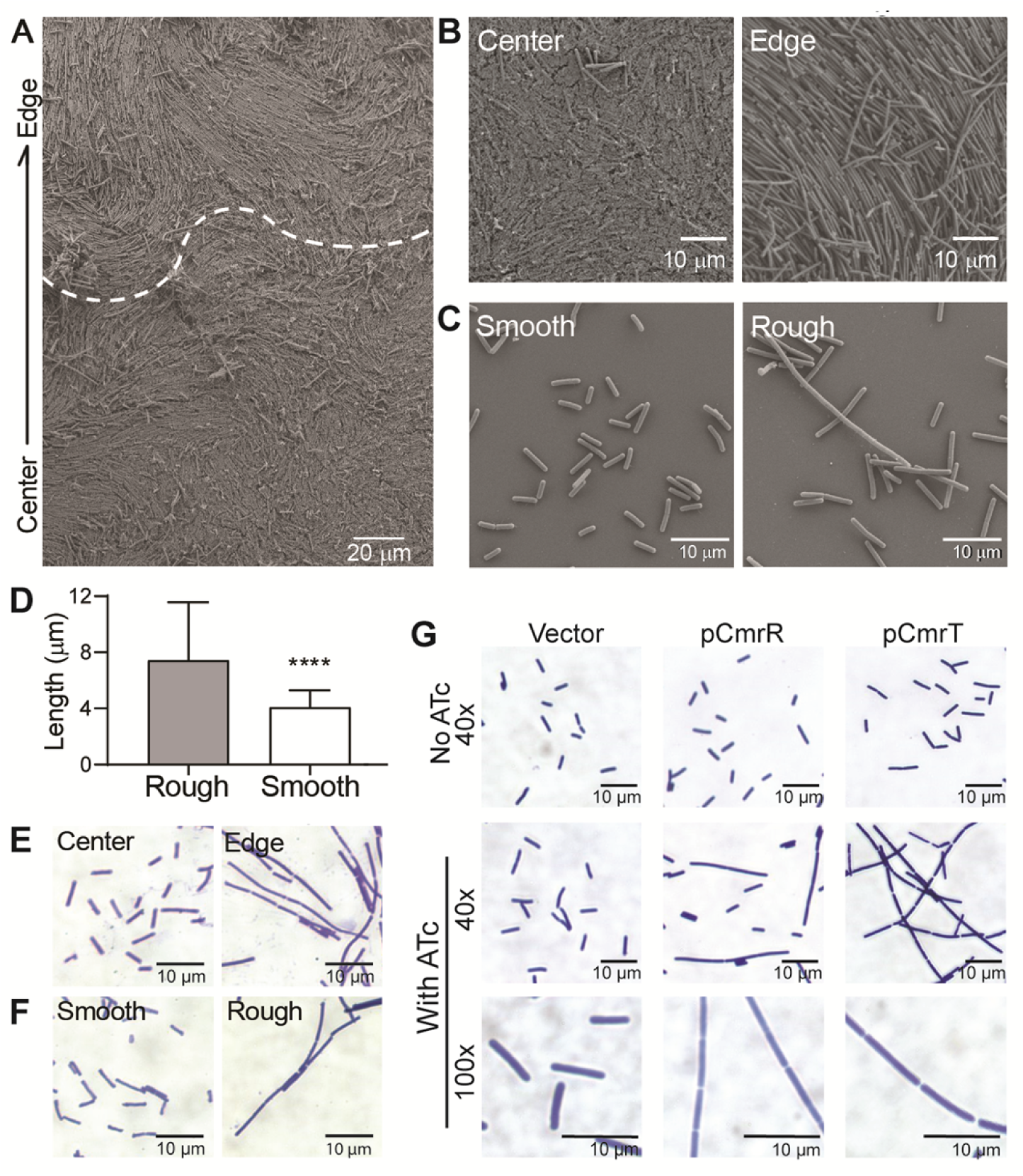
CmrR and CmrT promote bacterial chaining. A) SEM images of a whole R20291 colony after 3 days of growth on BHIS agar. (A) A colony excised on an agar slice, at 1 × 10^3^ magnification. The arrow denotes the orientation of the image with respect to the colony. The dotted line marks the apparent transition from the colony edge and center. (B) 2.5 × 10^3^ magnification images of cells at the colony center and edge. (C) 4.5 × 10^3^ magnification of rough and smooth cell suspensions. Samples were grown in broth culture prior to fixation. Shown are representative images from two biological replicates. (D) Quantification of cell lengths obtained from SEM images using ImageJ. Cells were measured from at least seven images from two biological replicates (rough N = 574, smooth N = 457). (E, F) Gram stain of *C. difficile* samples taken directly from the center and edge of a colony (E) or from rough and smooth colonies (F) at 60X magnification. G) Gram stain of *C. difficile* with plasmids for expression of *cmrR* or *cmrT*, or a vector control were grown on BHIS-agar with and without inducer (10 ng/mL ATc for vector and pCmrR; 2 ng/mL ATc for pCmrT). Magnifications are indicated. Shown are representative images from two independent experiments.

Bacterial motility, cell morphology, and adherent behaviors are central to biofilm development, and chaining and filamentation can also affect surface adhesion and biofilm formation (52–55). We therefore examined the role of CmrRST in *C. difficile* biofilm formation. Bacteria isolated from rough and smooth colonies and the *cmrR* and *cmrT* mutants were assayed for biofilm production in rich medium on plastic as described previously (36, 37). The *cmrR* mutant showed significantly increased biofilm formation compared to the rough isolate, while the *cmrT* mutant showed no significant difference (Figure S5), indicating that CmrR negatively regulates biofilm formation.

### Phase variable colony morphology via CmrRST impacts *C. difficile* virulence

Given the broad role of CmrRST in controlling *C. difficile* physiology and behaviors, we predicted that the *cmrR* or *cmrT* mutants would have altered virulence in a hamster model of *C. difficile* infection. Male and female Syrian golden hamsters were inoculated with spores of Δ*cmrR* or Δ*cmrT*, or smooth or rough isolates of R20291, which remain capable of phase varying *cmrRST* expression. Notably, the strains tested did not differ in sporulation or germination rates *in vitro* (Figure S6). The animals were monitored for disease symptoms and euthanized when moribund as described in the Materials and Methods. Hamsters infected with Δ*cmrR* showed increased survival compared to those infected with the rough isolate of the R20291 parent (*P* = 0.003, log-rank test). Survival of hamsters infected with Δ*cmrR* was also greater than that of animals infected with the smooth isolate, but the difference did not reach statistical significance (*P* = 0.098, log-rank test). While the mean times to morbidity were comparable for the animals that did succumb (44.08 hours for Δ*cmrR*, 48.15 ± 12.87 hours for rough, 54.69 ± 22.90 hours for smooth), most of the animals inoculated with Δ*cmrR* survived (Figure 7A). For animals infected with Δ*cmrT*, both time to morbidity and percent animal survival were equivalent to those infected with the rough and smooth isolates.

**Figure 7.**
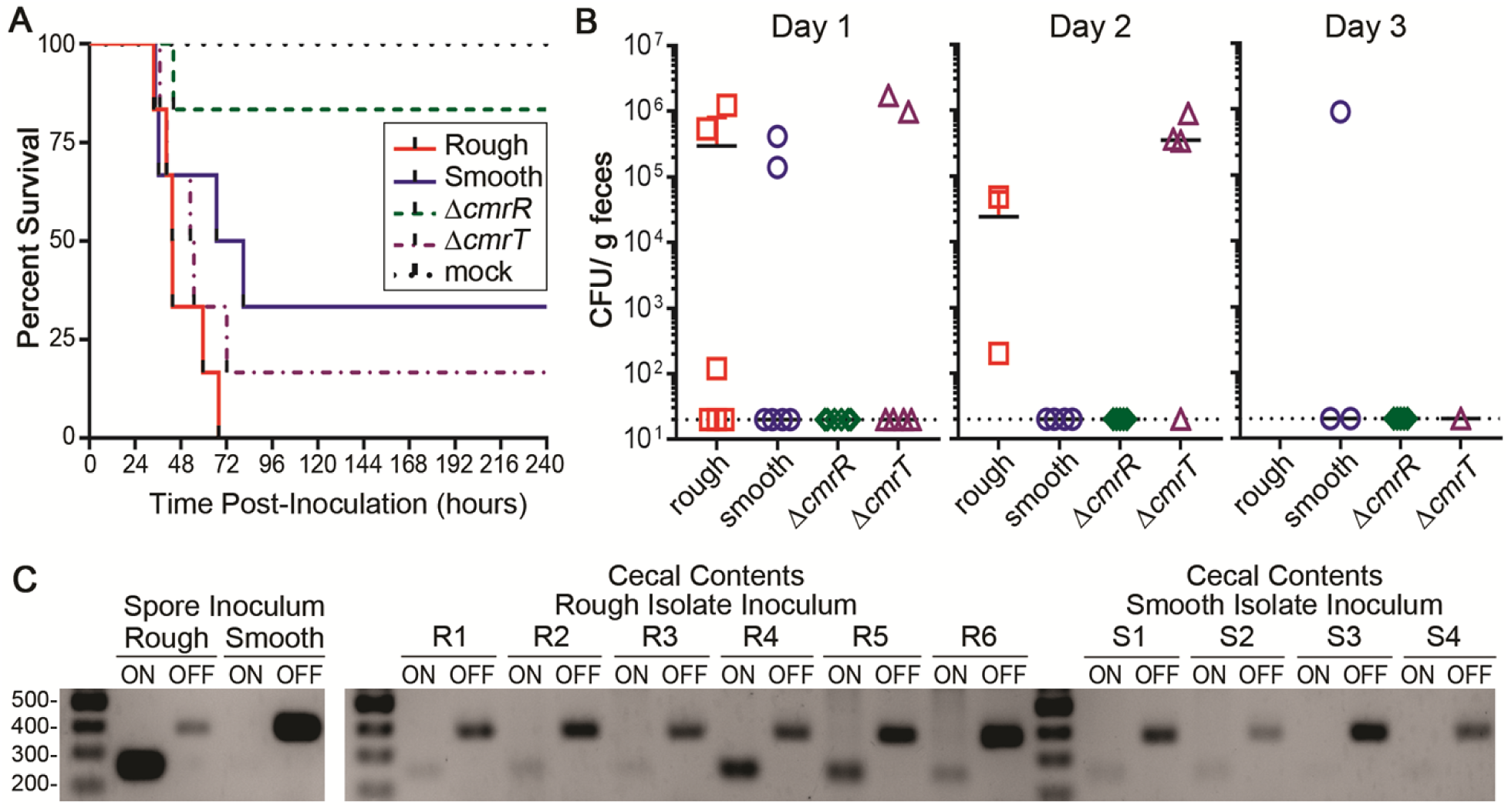
Colony morphology and CmrR impact *C. difficile* virulence. Male and female Syrian golden hamsters (N = 6) with were inoculated with 5000 spores of the indicated *C. difficile* strains/isolates. (A) Kaplan-Meier survival data showing time of morbidity and euthanasia. Log-rank test: rough v. Δ*cmrR*, *P* = 0.003; smooth v. Δ*cmrR, P* = 0.098, log-rank test. (B) CFU enumerated in feces collected at 24 hour intervals post-inoculation. Symbols indicate CFU from individual animals, and bars indicate the means. (C) Orientation of the *cmr* switch determined by PCR for the spore inoculums and cecal contents of moribund hamsters (rough R1-6, smooth S1-4) with detectable *C. difficile*.

To evaluate intestinal colonization, fecal samples were collected every 24 hours post-inoculation, and dilutions were plated on TCCFA medium (containing the spore germinant taurocholic acid) to enumerate *C. difficile* colony forming units (CFU). All animals infected with Δ*cmrT*, rough, and smooth isolates yielded detectable CFU at the time of collection preceding disease development and euthanasia (Figure 7B). However, we observed no correlation between bacterial load and disease onset or severity. The number of Δ*cmrR* CFU was never above the limit of detection in feces of infected animals (Figure 7B), though *C. difficile* was detected in the cecum of the Δ*cmrR*-infected hamster that became moribund (data not shown). These data indicate that CmrR, but not CmrT, is important for the ability of *C. difficile* to colonize the intestinal tract and cause disease in the hamster model. Importantly, the mutants do not produce lower levels of TcdA toxin than rough and smooth isolates *in vitro*, suggesting that the virulence defect of Δ*cmrR* occurs through a toxin-independent mechanism (Figure S6).

Based on this result, we would have expected the smooth isolate (CmrRST “off”) to be attenuated compared to the rough isolate (CmrRST “on”). While hamsters infected with the rough isolate all succumbed to disease, 33% of hamsters inoculated with the smooth isolate survived, suggesting moderately lower virulence of the smooth isolate (Figure 7A). However, this difference did not reach statistical significance as the survival times of hamsters that became moribund were not significantly different (P = 0.112, log-rank test), and both isolates were able to colonize (Figure 7A, B).

We considered the possibility that the rough and smooth isolates, though chosen based on distinct colony morphologies, are “wild-type” and capable of phase variation. Accordingly, selective pressures in the intestinal tract may have resulted in a phenotypic switch during infection, mitigating the observed differences in virulence. To test this, we determined the orientations of the *cmrRST* switch in the spore preparations and the contents of the cecums collected from infected animals immediately after sacrifice. DNA obtained from cecal contents was subjected to PCR with two primer sets that amplify from the *cmrRST* switch in a particular orientation. Analysis of the *cmrRST* switch orientation in the spore inoculums indicate that the rough isolate consisted of bacteria with the switch primarily in the ON orientation, while the smooth isolate contained those primarily in the OFF orientation, consistent with prior *in vitro* analyses (Figure 7C). In the cecal contents of hamsters inoculated with the smooth isolate, *C. difficile* retained the *cmrRST* switch predominantly in the OFF orientation. In contrast, cecal contents from rough isolate-infected animals also contained the switch in a mixed or predominantly OFF orientation, indicating that that the rough/ON isolate inoculum underwent phase variation during infection. These data suggest that either ON/OFF switch rates are altered in favor of the OFF orientation during acute, short-term infection, and/or there is a selective pressure for the *cmrRST* OFF state and corresponding phenotypes.

## Discussion

Phase variation provides a mechanism for generating phenotypic heterogeneity in bacterial populations to enhance the survival of the population as a whole when encountering detrimental environmental stresses. In this study, we characterized the phase variation of a signal transduction system, CmrRST, which we found broadly impacts *C. difficile* physiology and behavior. During growth *in vitro*, *C. difficile* populations consist of bacterial subpopulations that are CmrRST ON, yielding rough colonies, and CmrRST OFF, yielding smooth colonies. These subtypes display divergent motility phenotypes *in vitro*, with smooth colony isolates showing increased swimming motility and rough colony isolates displaying enhanced surface motility. The identification of growth conditions that segregate for the respective morphological variants, each with enhanced motility in the corresponding growth condition, indicates a selective advantage for that motility behavior.

Although ectopic expression of either response regulator, CmrR or CmrT, increased surface motility, only CmrT is required for surface migration and rough colony development, suggesting that CmrT is the dominant regulator under these conditions. Alternatively, CmrR may also play a role but requires an activating signal through the histidine kinase CmrS to be functional. The latter possibility is supported by the observation that a phosphomimic substitution at the predicted phosphorylation site in the receiver domain increases CmrR activity. In contrast, because CmrT contains a glutamic acid at this site, it likely does not require activation by a histidine kinase and instead acts constitutively. For this reason, the receiver domain of CmrT may be considered pseudo-receiver domain (47, 56).

In the hamster model of acute infection, the *cmrR* mutant did not detectably colonize and exhibited reduced virulence, indicating a role for this signal transduction system during infection. In contrast with the *in vitro* motility phenotypes driven primarily by CmrT, the CmrR regulator appears to be critical for virulence. The divergent roles for the regulators suggest that an activating signal for CmrR is present in the host intestine, though it remains possible that CmrT activity is somehow inhibited during infection. A similar mechanistic distinction between the regulators may explain the role of CmrR, but not CmrT, in biofilm formation *in vitro*. Whether CmrR and CmrT control distinct regulons, act cooperatively, and/or antagonistically is unknown and currently under investigation.

Compared to the strong attenuated virulence of the *cmrR* mutant, the rough and smooth isolates exhibited modest differences in virulence. The latter result may have arisen from the ability of the isolates to phase vary, resulting in convergent phenotypes due to selective pressures in the intestinal environment. Tracking of the *cmr* switch orientations in the inoculums and cecal contents at the experimental endpoint indicate that the rough variant (*cmrRST* ON) switched to populations with the *cmr* switch in a mixed or predominantly OFF orientation. That phase variation of CmrRST occurred during infection may have resulted in diminished differences in virulence between the rough and smooth variants. In addition, these findings imply the presence of a selective pressure that favors the *cmrRST* OFF phase variant, at least in the later stages of infection. However, the high conservation of the *cmrRST* locus and the upstream regulatory region indicates that this regulatory system and its ability to phase vary confer a fitness advantage. The colonization defect of the *cmrR* mutant suggests that signaling through CmrR contributes to initial colonization, and therefore the ability to generate a CmrRST ON subpopulation may be important at an early stage of infection. In our study, cecal contents were harvested at the experimental endpoint when disease was fulminant, and the collection of samples after more frequent intervals is limited by the short duration of the infection and low bacterial load prior to 24 hours post-inoculation. This sampling strategy may not capture transient changes in the *cmr* switch bias during different stages of infection, over a longer infection, or in a different location in the intestinal tract. Testing the impact of *cmrR* and *cmrT* mutations in a phase-locked ON background as well as a double *cmrR cmrT* mutant may better reveal the role of this system in *C. difficile* virulence.

Unlike phase variation mechanisms that control the production of a specific surface factor, the phase variation of the CmrRST system is poised to coordinate the expression of multiple pathways. Previous studies have described phase variation of proteins with regulatory function resulting in broad changes in gene expression. A few transcriptional regulators have been reported to phase vary, including the global virulence gene regulators BvgAS in *Bordetella* and the response regulator FlgR in *Campylobacter jejuni* (57–59). In *C. difficile*, phase variation of flagellar genes impacts the transcription of the alternative sigma factor gene *sigD*, which couples the expression of flagellum and toxin genes, linking virulence factor production to the orientation of the flagellar switch (19, 60). Coordinated regulation of genes resulting in global changes in transcription can also result from phase variation of DNA modification systems, as observed in *Haemophilus influenza*, *Helicobacter pylori*, *Salmonella enterica*, and *Streptococcus strains.* (61–66). In *Streptococcus pneumoniae*, for example, DNA inversions within three genes encoding the sequence recognition factor of a Type I restriction-modification (RM) system allow for changes in genomic methylation patterns, resulting in phase variable regulation of virulence factors including capsule (66, 67). The CmrR and CmrT response regulators each contain DNA binding domains and have the capacity to regulate gene transcription. Therefore, inversion of the *cmr* switch likely also indirectly controls the expression of multiple genes in a phase variable manner.

The CmrRST system regulates multiple processes, but the basis for the phenotypic changes are unclear. Colony morphology and motility assays indicate TFP- and flagellum-independent mechanisms, and the transcription of TFP and flagellum genes is not affected by CmrR and CmrT. TFP-independent surface migration has not previously been described in *C. difficile*. Examination of cellular morphology indicates that CmrRST promotes bacterial chaining, which may result in bacterial migration as cells are pushed along the axis of growth and division, as observed in *Bacillus subtilis* (68). This mechanism appears similar to gliding motility in *Clostridium perfringens*, although that process requires TFP (69, 70). Bacterial chaining is observed in a number of bacterial species and can affect flagellar motility, surface adhesion, biofilm formation, susceptibility to phagocytosis, and virulence (51-55, 68, 71-74). Chaining of *C. difficile* through regulation by CmrRST may similarly alter motility behaviors and persistence in the intestine. How CmrRST mediates changes to cell chaining, colony morphology, and motility is unknown, and ongoing work to define the CmrRST regulon may reveal the underlying mechanisms.

The complexity of *cmrRST* regulation not only alludes to the importance of controlling CmrRST activity, but also highlights the growing link between phase variation and c-di-GMP signaling in *C. difficile*. Expression of flagellar genes is also modulated by both a c-di-GMP binding riboswitch and an invertible element (19, 40, 43), as well as two additional phase variable genes in *C. difficile* that encode c-di-GMP phosphodiesterases (28). In the case of flagellar gene regulation, c-di-GMP inhibits transcription through a riboswitch that induces transcription termination, and the invertible flagellar switch further controls expression through an uncharacterized post-transcriptional mechanism. For *cmrRST* expression, c-di-GMP positively regulates transcription, and the *cmr* switch controls phase variation through an undefined mechanism (28, 39). The hierarchy between these regulatory elements is unclear. A σ^A^-dependent promoter lies 5’ of both the riboswitch and *cmr* switch (41). The invertible sequence may contain an additional promoter to drive expression when properly oriented (4, 75). Alternatively, the upstream σ^A^-dependent promoter may be the only site of transcriptional initiation, and the invertible element may contain another regulatory sequence that modulates expression (76).

Finally, this and other recent work underscores that genetically clonal *C. difficile* strains, as well as isogenic mutants, may differ phenotypically due to phase variation. The seven sites of site-specific DNA recombination in the *C. difficile* genome can be inverted independently, and this study shows that *in vitro* culturing conditions can change the biases of switch orientations in the overall population. From a research standpoint, this can introduce significant variability into otherwise controlled experiments. This concern may be especially significant in infection studies, where differences in switch biases in inoculums may skew apparent outcomes. Caution should be taken in interpreting such data, as observed differences may be due to differences in expression of phase variable traits rather than the targeted mutations.

Through phase variation, populations of bacterial pathogens can rapidly adapt to changing environmental pressures, balancing the need to move, adhere, and avoid immune recognition through the course of infection. The phase variation of CmrRST may allow alternating modes of surface or swimming motility in *C. difficile* as needed during infection –swimming motility (*cmrRST* OFF) may allow exploration and colonization of new sites, while bacterial chaining (*cmrRST* ON) may allow the spread of bacteria along the epithelial surface once contact has been made. Further work understanding the role of CmrRST will provide important insights into *C. difficile* pathogenesis.

## Methods

### Growth and maintenance of bacterial strains

*C. difficile* strains were maintained in an anaerobic environment of 85% N_2_, 5% CO_2_, and 10% H_2_. *C. difficile* strains were cultured in Tryptone Yeast (TY; 30 g/liter Bacto tryptone, 20 g/liter yeast extract, 1 g/liter thioglycolate) broth or Brain Heart Infusion plus yeast (BHIS; 37 g/liter Bacto brain heart infusion, 5 g/liter yeast extract) media at 37 °C with static growth. *Escherichia coli* strains were grown in Luria Bertani medium at 37 °C with aeration. Antibiotics were used where indicated at the following concentrations: chloramphenicol (Cm), 10 µg/mL; thiamphenicol (Tm), 10 µg/mL; kanamycin (Kan), 100 µg/mL; ampicillin (Amp), 100 µg/mL. Table S1 lists descriptions of the strains and plasmids used in this study.

### Differentiation of rough and smooth colonies

To differentiate rough and smooth colonies from wild-type *C. difficile* strains, 5 µL of liquid cultures were spotted onto BHIS agar plates and grown for 48-72 hours. For rough and smooth colonies, all growth was collected and plated in dilutions on BHIS agar. For predominantly rough colonies, growth was collected from the filamentous edges (18, 33). For predominantly smooth colonies, growth was collected from the center of the spot (our stock of *C. difficile* R20291 consists primarily of bacteria yielding smooth colonies, so the spot center represents the inoculum). For enumeration of rough and smooth colonies, serial dilutions were plated, rough and total colony forming units (CFU) were counted, and data were expressed as percent rough CFU.

### Microscopy

For whole colony imaging, colonies were grown on BHIS plates for 24 hours prior to imaging. For strains containing plasmids, the BHIS agar medium contained 10 µg/mL Tm. For strains carrying pDccA, pDccA^mut^, pEAL, or pMC-Pcpr, 2 µg/mL nisin was included to induce gene expression. For strains with pCmrR, pCmrT, or mutant derivatives, anhydrotetracycline (ATc) was added at the indicated concentrations. Colonies were imaged at 20-80x magnification using a Zeiss Stereo Discovery V8 dissecting microscope with a glass stage and light from the top.

For Gram stain imaging, rough or smooth colonies were differentiated as previously described and heat fixed on a glass slide. Cells were Gram-stained (BD Kit 212524) and visualized at 40-80x magnification using an Olympus BX60 compound microscope.

For visualization by scanning electron microscopy, R20291 rough and smooth isolates and the *sigD* mutant were grown overnight in TY broth and washed once with phosphate buffered saline (PBS). For visualization of colony structure, R20291 was spotted on BHIS agar and grown for 3 days. The colony was excised on agar and fixed. All samples were fixed in 4% glutaraldehyde in 150 mM sodium phosphate buffer. The samples were dehydrated through increasing concentrations of ethanol and dried using carbon dioxide as the transition solvent in a Samdri-795 critical-point dryer (Tousimis Research Corporation, Rockville, MD). Coverslips were mounted on aluminum stubs with carbon adhesive tabs, followed by a 5-nm thickness platinum sputter in a Hummer X sputter coater (Anatech USA, Union City, CA). Images were taken with a working distance of 5 mm and a 130-μm aperture using a Zeiss Supra 25 field emission scanning electron microscope (FESEM) operating at 5 kV (Carl Zeiss SMT Inc., Peabody, MA). For quantification of cell length, at least seven images from two biological replicates were analyzed using ImageJ.

### Quantitative PCR analysis of invertible element orientations

Quantitative PCR (qPCR) was used to quantify the percent of the population with each of the seven invertible elements in a given orientation as described previously (28). Rough and smooth colonies were collected from a BHIS plate, and genomic DNA was purified. qPCR was performed with SensiMix SYBR (BioLine). Twenty μL reactions with 100 ng of genomic DNA and 100 nM primers were used. Reactions were run on Lightcycler 96 system (Roche) with the following three-step cycling conditions: 98°C for 2 min, followed by 40 cycles of 98°C for 30 s, 60°C for 1 min, and 72°C for 30 sec. Quantification was done using the ΔΔ*C_T_* method, with *rpoA* as the reference gene and the indicated reference condition. All primers used in this and other experiments are listed in Table S2.

### Construction of Δ*cmrR* and Δ*cmrT* in *C. difficile* R20291

Markerless deletions of *cmrR* (locus tag CDR20291_3128) and *cmrT* (locus tag CDR20291_3126) were done using a previously described *codA*-based allelic exchange method with minor modifications (77). Briefly, approximately 1100-1300 bp genomic fragments were PCR-amplified upstream and downstream of *cmrR* and *cmrT* with the following primers sets: OS158 and OS266 (*cmrR*, upstream), OS163 and OS267 (*cmrR*, downstream); OS268 and OS269 (*cmrT*, upstream), OS271 and OS283 (*cmrT*, downstream). Complementary overlapping sequences were added to the 5-prime end of primers to allow for accurate fusion of all PCR products into PmeI-linearized pMTL-SC7215 vector using Gibson Assembly Master Mix (New England BioLabs). The resulting plasmids were then conjugated into *C. difficile* R20291 strain (GenBank accession: CBE06969.1) using the heat-stimulated conjugation method described elsewhere (78). Mutants were selected as previously described and screened by colony-PCR for the presence of the correct left and right homology-chromosomal junction and the absence of respective *cmrR* and *cmrT* coding sequence (77). All PCR products were Sanger-sequenced to confirm the correct genetic construct and the absence of any secondary mutations.

### Surface and swimming motility assays

For surface motility assays, 5 µL of overnight (16-18hr) cultures were spotted onto BHIS-1.8% agar supplemented with 1% glucose (38). For swimming motility assays, 1 µL of overnight cultures was inoculated into 0.5x BHIS-0.3% agar (19). For plasmid-bearing strains in both assays, the medium was supplemented with 10 µg/mL Tm. Where indicated, ATc was included at the indicated concentrations to induce expression. At 24 hour intervals the diameters of growth were taken as the average of two perpendicular measurements. Data from at least 8 biologically independent cultures were analyzed using a one-way ANOVA to determine statistical significance. The plates were photographed using a Syngene G:Box imager.

### Biofilm Assay

Biofilm assays were done as previously described (37). Overnight cultures of *C. difficile* were diluted 1:100 in BHIS 1% glucose 50 mM sodium phosphate buffer (pH 7.5) in 24-well polystyrene plates. After 24 hours of growth, supernatants were removed, the biofilms were washed once with PBS and then stained for 30 minutes with 0.1% crystal violet. After 30 minutes, the biofilms were washed again with PBS, and the crystal violet was solubilized with ethanol. Absorbance was read at 570 nm. Four independent experiments were performed.

### RNA isolation and real-time PCR

*C. difficile* with vector or *cmrR*/*cmrT* expression plasmids were grown for 48 hours in 0.5x BHIS-0.3% agar to express flagellar genes. Bacteria were recovered and cultured in TY broth with 10 ng/mL ATc for vector and pCmrR; 2 ng/mL ATc for pCmrT. Samples were collected at mid-exponential phase and preserved in 1:1 ethanol-acetate. RNA was isolated, treated with DNase I, and reverse transcribed including a no-reverse transcriptase control as previously described (31, 33). Real-time PCRs were done using 2 ng of cDNA, primers at a final concentration of 500 nM, and SYBR Green Real-Time qPCR reagents (Thermo Fisher) with an annealing temperature of 55°C. The data were analyzed using the ΔΔCt method with *rpoC* as the reference gene and normalized to the uninduced condition.

### Animal experiments

All animal experimentation was performed under the guidance of veterinarians and trained animal technicians within the Emory University Division of Animal Resources (DAR). Animal experiments were performed with prior approval from the Emory University Institutional Animal Care and Use Committee (IACUC). *C. difficile* spores were collected from 70:30 sporulation broth after 3 days of growth (79). The spores were purified using a sucrose gradient and stored in PBS with 1% bovine serum albumin as described previously (44, 80). Prior to inoculation, the spores were enumerated by plating serial dilutions on BHIS-agar containing 0.1% sodium taurocholate.

Male and female Syrian golden hamsters (*Mesocricetus auratus*, strain LVG, 7-8 weeks old, Charles River Laboratories) were housed individually in sterile cages and given a standard rodent diet and water *ad libitum*. To render the animals susceptible to *C. difficile*, one dose of clindamycin (30 mg kg^−1^ of body weight) was administered by oral gavage seven days prior to inoculation. Hamsters were inoculated with ∼5000 spores of a single strain of *C. difficile* (80, 81). Uninfected controls treated with clindamycin were included in each experiment. Hamsters were weighed at least daily and monitored for signs of disease (weight loss, lethargy, diarrhea and wet tail). Fecal samples were collected daily for determination of bacterial burden (see below). Hamsters were considered moribund if they lost 15% or more of their highest weight or if they showed disease symptoms of lethargy, diarrhea and wet tail. Animals meeting either criterion were euthanized by CO_2_ asphyxiation and thoracotomy. Immediately following euthanasia, animals were necropsied, and cecal contents were obtained for enumeration of total *C. difficile* CFU and for subsequent DNA isolation. To enumerate *C. difficile* CFU, fecal and cecal samples were weighed, suspended in 1X PBS, heated to 60°C for 20 min to minimize growth of other organisms, and plated on taurocholate cycloserine cefoxitin fructose agar (TCCFA; (82, 83)). *C. difficile* CFU were enumerated after 48 hours. Six animals (3 male, 3 female) per *C. difficile* strain were tested, and the data were analyzed using GraphPad Prism.

### Semi-quantitative PCR analysis of DNA from hamster ceca

Hamster cecal contents in PBS were treated with lysozyme and subjected to bead beating to lyse spores. DNA was purified by phenol:chloroform:isopropanol extraction and washed with ethanol. Subsequently, 0.5 ng DNA per 50 µL reaction was PCR amplified using 0.5 μM orientation specific primers to detect the orientation of the *cmr* switch (Cdi6): RT2378-RT2379 for the ON orientation; RT2378-RT2197 for the OFF orientation.

### Sporulation assays

*C. difficile* cultures were grown in BHIS medium supplemented with 0.1% taurocholate and 0.2% fructose until mid-exponential phase (i.e., OD_600_ of 0.5), and 0.25 ml aliquots were plated onto 70:30 agar supplemented with 2 µg/ml thiamphenicol (79). After 24 h growth, ethanol resistance assays were performed as previously described (84, 85). Briefly, the cells were removed from plates after 24 h (H_24_) and suspended in BHIS medium to an OD_600_ of ∼1.0. The number of vegetative cells per milliliter was determined by immediately serially diluting and plating the suspended cells onto BHIS. Simultaneously, a 0.5-ml aliquot was added to 0.3 ml of 95% ethanol and 0.2 ml of dH_2_O to achieve a final concentration of 28.5% ethanol, vortexed, and incubated for 15 min to eliminate all vegetative cells; ethanol-treated cells were subsequently serially diluted in 1× PBS with 0.1% taurocholate and applied to BHIS plus 0.1% taurocholate plates to determine the total number of spores. After at least 24 h of growth, CFU were enumerated, and the sporulation frequency was calculated as the total number of spores divided by the total number of viable cells (spores plus vegetative cells). A nonsporulating mutant (MC310; *spo0A::erm*) was used as a negative control.

### Germination assays

*C. difficile* strains were grown in 500-ml liquid 70:30 medium and spores were purified for germination studies as previously described, with some modifications (81, 86–88). Cultures in sporulation medium were removed from the anaerobic chamber after 120 h of anaerobic growth and kept outside of the chamber in atmospheric oxygen overnight. Spore cultures were collected by centrifugation, suspended in water, and frozen for 15 min at −80°C. After thawing, spore suspensions were centrifuged for 10 min at ∼3200 × g in a swing bucket rotor, and the supernatant was discarded. Spore pellets were washed two times with water then suspended in 1 ml of a 1x PBS + 1% BSA solution, applied to a 12 ml 50% sucrose bed volume, and centrifuged at ∼3200 × g for 20 min. The supernatant was decanted and the spores were checked by phase-contrast microscopy for purity. Sucrose gradients were repeated until the preparations reached a purity of greater than 95%. Spore pellets were then washed three times with 1x PBS + 1% BSA and suspended to an OD_600_ = 3.0. Germination assays were carried out as previously described, with slight modifications (86, 87). After treatment of spores for 30 min at 60°C, spore germination was analyzed in BHIS containing 10 mM taurocholate and 100 mM glycine. The OD_600_ was determined immediately and at various time points during incubation at room temperature. Results are presented as means and standard errors of the means of three independent biological replicates.

### Detection of TcdA by western blot

Rough and smooth isolates as well as *cmrR* and *cmrT* mutants were grown for 24 hours in TY broth and normalized to OD_600_ ∼ 1.8 prior to collection. Cells were pelleted, suspended in SDS-PAGE buffer, and boiled for 10 minutes. Samples were then run on 4–20% Mini-PROTEAN TGX Precast Protein Gels (Bio Rad) and transferred to a nitrocellulose membrane. TcdA was detected as described previously using a mouse α-TcdA primary antibody (Novus Biologicals) and goat anti-mouse IgG conjugated with IR800 (Thermo Fisher) [42].

### Ethics Statement

All animal experimentation was performed under the guidance of veterinarians and trained animal technicians within the Emory University Division of Animal Resources (DAR). Animal experiments were performed with prior approval (approval number 201700396) from the Emory University Institutional Animal Care and Use Committee (IACUC) under protocol #DAR-2001737-052415BA. Animals considered moribund were euthanized by CO_2_ asphyxiation followed by thoracotomy in accordance with the Panel on Euthanasia of the American Veterinary Medical Association. The University is in compliance with state and federal Animal Welfare Acts, the standards and policies of the Public Health Service, including documents entitled “Guide for the Care and Use of Laboratory Animals” – National Academy Press, 2011, “Public Health Service Policy on Humane Care and Use of Laboratory Animals” – September 1986, and Public Law 89-544 with subsequent amendments. Emory University is registered with the United States Department of Agriculture (57-R-003) and has filed an Assurance of Compliance statement with the Office of Laboratory Animal Welfare of the National Institutes of Health (A3180-01).

## Acknowledgments

We thank the University of North Carolina (UNC) Microscopy Services Laboratory for assistance with scanning electron microscopy. The Microscopy Services Laboratory, Department of Pathology and Laboratory Medicine, is supported in part by P30 CA016086 Cancer Center Core Support Grant to the UNC Lineberger Comprehensive Cancer Center.

## References

1. Ryall B, Eydallin G, Ferenci T. Culture History and Population Heterogeneity as Determinants of Bacterial Adaptation: the Adaptomics of a Single Environmental Transition. Microbiol Mol Biol Rev. 2012;76(3):597–625.

2. Magdanova LA, Golyasnaya NVJM. Heterogeneity as an adaptive trait of microbial populations. Microbiol. 2013;82(1):1–10.

3. Casadesús J, Low DA. Programmed Heterogeneity: Epigenetic Mechanisms in Bacteria. J Biol Chem. 2013;288(20):13929–35.

4. van der Woude MW, Baumler AJ. Phase and antigenic variation in bacteria. Clin Microbiol Rev. 2004;17(3):581–611.

5. Grindley ND, Whiteson KL, Rice PA. Mechanisms of site-specific recombination. Annu Rev Biochem. 2006;75:567–605.

6. Weiser JN, Austrian R, Sreenivasan PK, Masure HR. Phase variation in pneumococcal opacity: relationship between colonial morphology and nasopharyngeal colonization. Infect Immun. 1994;62(6):2582–9.

7. Lim JK, Gunther NWt, Zhao H, Johnson DE, Keay SK, Mobley HL. In vivo phase variation of *Escherichia coli* type 1 fimbrial genes in women with urinary tract infection. Infect Immun. 1998;66(7):3303–10.

8. Li X, Lockatell CV, Johnson DE, Mobley HL. Identification of MrpI as the sole recombinase that regulates the phase variation of MR/P fimbria, a bladder colonization factor of uropathogenic *Proteus mirabilis*. J Molec Microbiol. 2002;45(3):865–74.

9. Briles DE, Novak L, Hotomi M, van Ginkel FW, King J. Nasal colonization with *Streptococcus pneumoniae* includes subpopulations of surface and invasive pneumococci. Infect Immun. 2005;73(10):6945–51.

10. Tauseef I, Ali YM, Bayliss CD. Phase variation of PorA, a major outer membrane protein, mediates escape of bactericidal antibodies by *Neisseria meningitidis*. Infect Immun. 2013;81(4):1374–80.

11. Alamro M, Bidmos FA, Chan H, Oldfield NJ, Newton E, Bai X, et al. Phase variation mediates reductions in expression of surface proteins during persistent meningococcal carriage. Infect Immun. 2014;82(6):2472–84.

12. Oliver MB, Roy AB, Kumar R, Lefkowitz EJ, Swords WE. *Streptococcus pneumoniae* TIGR4 phase-locked opacity variants differ in virulence phenotypes. mSphere. 2017;2(6):e00386–17.

13. Arai J, Hotomi M, Hollingshead SK, Ueno Y, Briles DE, Yamanaka N. *Streptococcus pneumoniae* isolates from middle ear fluid and nasopharynx of children with acute otitis media exhibit phase variation. J Clin Microbiol. 2011;49(4):1646–9.

14. Kuehne SA, Cartman ST, Heap JT, Kelly ML, Cockayne A, Minton NP. The role of toxin A and toxin B in *Clostridium difficile* infection. Nature. 2010;467(7316):711–3.

15. Shen A. *Clostridium difficile* toxins: mediators of inflammation. J Innate Immun. 2012;4(2):149–58.

16. Carter GP, Chakravorty A, Pham Nguyen TA, Mileto S, Schreiber F, Li L, et al. Defining the Roles of TcdA and TcdB in Localized Gastrointestinal Disease, Systemic Organ Damage, and the Host Response during *Clostridium difficile* Infections. mBio. 2015;6(3):e00551–15.

17. Morris GN, Winter J, Cato EP, Ritchie AE, Bokkenheuser VD. *Clostridium scindens* sp. nov., a human intestinal bacterium with desmolytic activity on corticoids. Int J Systematic Bacteriol. 1985;35(4):478–81.

18. Lipovsek S, Leitinger G, Rupnik M. Ultrastructure of *Clostridium difficile* colonies. Anaerobe. 2013;24:66–70.

19. Anjuwon-Foster BR, Tamayo R. A genetic switch controls the production of flagella and toxins in *Clostridium difficile*. PLoS Genet. 2017;13(3):e1006701.

20. Anjuwon-Foster BR, Tamayo R. Phase variation of *Clostridium difficile* virulence factors. Gut Microbes. 2018;9(1):76–83.

21. Simpson LM, White VK, Zane SF, Oliver JD. Correlation between virulence and colony morphology in *Vibrio vulnificus*. Infect Immun. 1987;55(1):269–72.

22. Kansal RG, Gomez-Flores R, Mehta RT. Change in colony morphology influences the virulence as well as the biochemical properties of the *Mycobacterium avium* complex. J Microbial Pathogenesis. 1998;25(4):203–14.

23. Lenz LL, Portnoy DA. Identification of a second Listeria secA gene associated with protein secretion and the rough phenotype. Molec Microbiol. 2002;45(4):1043–56.

24. Tipton KA, Dimitrova D, Rather PN. Phase-Variable Control of Multiple Phenotypes in *Acinetobacter baumannii* Strain AB5075. J Bacteriol. 2015;197(15):2593–9.

25. Clary G, Sasindran SJ, Nesbitt N, Mason L, Cole S, Azad A, et al. *Mycobacterium abscessus* smooth and rough morphotypes form antimicrobial-tolerant biofilm phenotypes but are killed by acetic acid. Antimicrob Agents Chemother. 2018;62(3):e01782–17.

26. Chin CY, Tipton KA, Farokhyfar M, Burd EM, Weiss DS, Rather PN. A high-frequency phenotypic switch links bacterial virulence and environmental survival in *Acinetobacter baumannii*. Nature Microbiol. 2018;3(5):563.

27. Dennis EA, Coats MT, Griffin S, Pang B, Briles DE, Crain MJ, et al. Hyperencapsulated mucoid pneumococcal isolates from patients with cystic fibrosis have increased biofilm density and persistence in vivo. Pathogens and Disease. 2018;76(7):fty073.

28. Sekulovic O, Mathias Garrett E, Bourgeois J, Tamayo R, Shen A, Camilli A. Genome-wide detection of conservative site-specific recombination in bacteria. PLoS Genet. 2018;14(4):e1007332.

29. Sekulovic O, Ospina Bedoya M, Fivian-Hughes AS, Fairweather NF, Fortier LC. The *Clostridium difficile* cell wall protein CwpV confers phase-variable phage resistance. Molec Microbiol. 2015;98(2):329–42.

30. Sekulovic O, Fortier LC. Global transcriptional response of *Clostridium difficile* carrying the CD38 prophage. Appl Environ Microbiol. 2015;81(4):1364–74.

31. Anjuwon-Foster BR, Maldonado-Vazquez N, Tamayo R. Characterization of flagellar and toxin phase variation in *Clostridiodes difficile* ribotype 012 isolates. J Bacteriol. 2018. 200(14):pii:e00056–18.

32. Aubry A, Hussack G, Chen W, Kuolee R, Twine SM, Fulton KM, et al. Modulation of toxin production by the flagellar regulon in *Clostridium difficile*. Infect Immun. 2012;80(10):3521–32.

33. Purcell EB, McKee RW, McBride SM, Waters CM, Tamayo R. Cyclic diguanylate inversely regulates motility and aggregation in *Clostridium difficile*. J Bacteriol. 2012;194(13):3307–16.

34. El Meouche I, Peltier J, Monot M, Soutourina O, Pestel-Caron M, Dupuy B, et al. Characterization of the SigD regulon of *C. difficile* and its positive control of toxin production through the regulation of *tcdR*. PloS One. 2013;8(12):e83748.

35. Bordeleau E, Purcell EB, Lafontaine DA, Fortier LC, Tamayo R, Burrus V. Cyclic di-GMP riboswitch-regulated type IV pili contribute to aggregation of *Clostridium difficile*. J Bacteriol. 2015;197(5):819–32.

36. Purcell EB, McKee RW, Bordeleau E, Burrus V, Tamayo R. Regulation of Type IV Pili Contributes to Surface Behaviors of Historical and Epidemic Strains of *Clostridium difficile*. J Bacteriol. 2015;198(3):565–77.

37. Purcell EB, McKee RW, Courson DS, Garrett EM, McBride SM, Cheney RE, et al. A nutrient-regulated cyclic diguanylate phosphodiesterase controls *Clostridium difficile* biofilm and toxin production during stationary phase. Infect Immun. 2017;85(9):pii: e00347–17.

38. Purcell EB, Tamayo R. Cyclic diguanylate signaling in Gram-positive bacteria. FEMS Microbiol Rev. 2016;40(5):753–73.

39. McKee RW, Harvest CK, Tamayo R. Cyclic Diguanylate Regulates Virulence Factor Genes via Multiple Riboswitches in *Clostridium difficile*. mSphere. 2018;3(5):e00423–18.

40. Sudarsan N, Lee ER, Weinberg Z, Moy RH, Kim JN, Link KH, et al. Riboswitches in eubacteria sense the second messenger cyclic di-GMP. Science (New York, NY). 2008;321(5887):411–3.

41. Soutourina OA, Monot M, Boudry P, Saujet L, Pichon C, Sismeiro O, et al. Genome-wide identification of regulatory RNAs in the human pathogen *Clostridium difficile*. PLoS Genet. 2013;9(5):e1003493.

42. Tamayo R. Cyclic diguanylate riboswitches control bacterial pathogenesis mechanisms. PLoS Pathog. 2019;15(2):e1007529.

43. McKee RW, Mangalea MR, Purcell EB, Borchardt EK, Tamayo R. The second messenger cyclic di-GMP regulates *Clostridium difficile* toxin production by controlling expression of *sigD*. J Bacteriol. 2013;195(22):5174–85.

44. McKee RW, Aleksanyan N, Garrett EM, Tamayo R. Type IV pili promote *Clostridium difficile* adherence and persistence in a mouse model of infection. Infect Immun. 2018;86(5):pii: e00943–17.

45. Peltier J, Shaw HA, Couchman EC, Dawson LF, Yu L, Choudhary JS, et al. Cyclic-di-GMP regulates production of sortase substrates of *Clostridium difficile* and their surface exposure through ZmpI protease-mediated cleavage. J Biol Chem. 2015;290(40):24453–69.

46. Lee ER, Baker JL, Weinberg Z, Sudarsan N, Breaker RR. An allosteric self-splicing ribozyme triggered by a bacterial second messenger. Science (New York, NY). 2010;329(5993):845–8.

47. Bourret RB. Receiver domain structure and function in response regulator proteins. Curr Opin Microbiol. 2010;13(2):142–9.

48. Lan C-Y, Igo MM. Differential expression of the OmpF and OmpC porin proteins in *Escherichia coli* K-12 depends upon the level of active OmpR. J Bacteriol. 1998;180(1):171–4.

49. Siam R, Marczynski GT. Glutamate at the phosphorylation site of response regulator CtrA provides essential activities without increasing DNA binding. Nucleic Acids Res. 2003;31(6):1775–9.

50. Kuhn M, Goebel W. Identification of an extracellular protein of *Listeria monocytogenes* possibly involved in intracellular uptake by mammalian cells. Infect Immun. 1989;57(1):55–61.

51. Rowan NJ, Candlish AA, Bubert A, Anderson JG, Kramer K, McLauchlin J. Virulent Rough Filaments of *Listeria monocytogenes* from Clinical and Food Samples Secreting Wild-Type Levels of Cell-Free p60 Protein. J Clin Microbiol. 2000;38(7):2643–8.

52. Young KD. The selective value of bacterial shape. Microbiol Mol Biol Rev. 2006;70(3):660–703.

53. Vejborg RM, Klemm P. Cellular chain formation in *Escherichia coli* biofilms. J Microbiol. 2009;155(5):1407–17.

54. Kan A, Del Valle I, Rudge T, Federici F, Haseloff J. Intercellular adhesion promotes clonal mixing in growing bacterial populations. J Royal Soc Interface. 2018;15(146):20180406.

55. Branda SS, González-Pastor JE, Ben-Yehuda S, Losick R, Kolter R. Fruiting body formation by *Bacillus subtilis*. Proc Natl Acad Sci U S A. 2001;98(20):11621–6.

56. Maule AF, Wright DP, Weiner JJ, Han L, Peterson FC, Volkman BF, et al. The aspartate-less receiver (ALR) domains: distribution, structure and function. PLoS Pathog 2015;11(4):e1004795.

57. Stibitz S, Aaronson W, Monack D, Falkow S. Phase variation in *Bordetella pertussis* by frameshift mutation in a gene for a novel two-component system. Nature.1989;338(6212):266–9.

58. Decker KB, James TD, Stibitz S, Hinton DM. The *Bordetella pertussis* model of exquisite gene control by the global transcription factor BvgA. J Microbiol. 2012;158(Pt 7):1665.

59. Hendrixson DR. A phase-variable mechanism controlling the *Campylobacter jejuni* FlgR response regulator influences commensalism. Mol Microbiol. 2006;61(6):1646–59.

60. Anjuwon-Foster BR, Tamayo R. Phase Variation of *Clostridium difficile* Virulence Factors. Gut Microbes. 2017:0.

61. De Bolle X, Bayliss CD, Field D, van de Ven T, Saunders NJ, Hood DW, et al. The length of a tetranucleotide repeat tract in *Haemophilus influenzae* determines the phase variation rate of a gene with homology to type III DNA methyltransferases. Mol Microbiol. 2000;35(1):211–22.

62. de Vries N, Duinsbergen D, Kuipers EJ, Pot RG, Wiesenekker P, Penn CW, et al. Transcriptional phase variation of a type III restriction-modification system in *Helicobacter pylori*. J Bacteriol. 2002;184(23):6615–23.

63. Srikhanta YN, Maguire TL, Stacey KJ, Grimmond SM, Jennings MP. The phasevarion: a genetic system controlling coordinated, random switching of expression of multiple genes. Proc Natl Acad Sci U S A. 2005;102(15):5547–51.

64. Broadbent SE, Davies MR, van der Woude MW. Phase variation controls expression of Salmonella lipopolysaccharide modification genes by a DNA methylation-dependent mechanism. Molec Microbiol. 2010;77(2):337–53.

65. Atack JM, Weinert LA, Tucker AW, Husna AU, Wileman TM, N FH, et al. *Streptococcus suis* contains multiple phase-variable methyltransferases that show a discrete lineage distribution. Nucleic Acids Res. 2018;46(21):11466–76.

66. Manso AS, Chai MH, Atack JM, Furi L, De Ste Croix M, Haigh R, et al. A random six-phase switch regulates pneumococcal virulence via global epigenetic changes. Nature Comm. 2014;5:5055.

67. Li J, Zhang JR. Phase Variation of *Streptococcus pneumoniae*. Microbiol Spectrum. 2019;7(1) doi: 10.1128/microbiolspec.GPP3-0005-2018.

68. van Gestel J, Vlamakis H, Kolter R. From cell differentiation to cell collectives: *Bacillus subtilis* uses division of labor to migrate. PLoS Biology. 2015;13(4):e1002141.

69. Varga JJ, Nguyen V, O’Brien DK, Rodgers K, Walker RA, Melville SB. Type IV pili-dependent gliding motility in the Gram-positive pathogen *Clostridium perfringens* and other Clostridia. Molec Microbiol. 2006;62(3):680–94.

70. Liu H, Bouillaut L, Sonenshein AL, Melville SB. Use of a mariner-based transposon mutagenesis system to isolate *Clostridium perfringens* mutants deficient in gliding motility. J Bacteriol. 2013;195(3):629–36.

71. Wuenscher MD, Köhler S, Bubert A, Gerike U, Goebel W. The *iap* gene of *Listeria monocytogenes* is essential for cell viability, and its gene product, p60, has bacteriolytic activity. J Bacteriol. 1993;175(11):3491–501.

72. Gioppo NM, Elias Jr WP, Vidotto MC, Linhares RE, Saridakis HO, Gomes TA, et al. Prevalence of HEp-2 cell-adherent *Escherichia coli* and characterisation of enteroaggregative *E. coli* and chain-like adherent *E. coli* isolated from children with and without diarrhoea, in Londrina, Brazil. FEMS Microbiol Lett. 2000;190(2):293–8.

73. Mercier C, Durrieu C, Briandet R, Domakova E, Tremblay J, Buist G, et al. Positive role of peptidoglycan breaks in lactococcal biofilm formation. Molec Microbiol. 2002;46(1):235–43.

74. Champion JA, Mitragotri S. Role of target geometry in phagocytosis. Proc Natl Acad Sci U S A. 2006;103(13):4930–4.

75. Schwan WR. Regulation of *fim* genes in uropathogenic *Escherichia coli*. World J Clin Infect Dis. 2011;1(1):17–25.

76. Emerson JE, Reynolds CB, Fagan RP, Shaw HA, Goulding D, Fairweather NF. A novel genetic switch controls phase variable expression of CwpV, a *Clostridium difficile* cell wall protein. Molec Microbiol. 2009;74(3):541–56.

77. Cartman ST, Kelly ML, Heeg D, Heap JT, Minton NP. Precise manipulation of the *Clostridium difficile* chromosome reveals a lack of association between the *tcdC* genotype and toxin production. Appl Environ Microbiol. 2012;78(13):4683–90.

78. Kirk JA, Fagan RP. Heat shock increases conjugation efficiency in *Clostridium difficile*. Anaerobe. 2016;42:1–5.

79. Putnam EE, Nock AM, Lawley TD, Shen A. SpoIVA and SipL are *Clostridium difficile* spore morphogenetic proteins. J Bacteriol. 2013;195(6):1214–25.

80. Edwards AN, Tamayo R, McBride SM. A novel regulator controls *Clostridium difficile* sporulation, motility and toxin production. Mol Microbiol. 2016;100(6):954–71.

81. Nawrocki KL, Edwards AN, Daou N, Bouillaut L, McBride SM. CodY-dependent regulation of sporulation in *Clostridium difficile*. J Bacteriol. 2016;198(15):2113–30.

82. George W, Sutter V, Citron D, Finegold S. Selective and differential medium for isolation of *Clostridium difficile*. J Clin Microbiol. 1979;9(2):214–9.

83. Wilson KH, Kennedy MJ, Fekety FR. Use of sodium taurocholate to enhance spore recovery on a medium selective for *Clostridium difficile*. J Clin Microbiol. 1982;15(3):443–6.

84. Edwards AN, McBride SM. Isolating and Purifying *Clostridium difficile* Spores. Methods Molec Biol. 2016;1476:117–28.

85. Childress KO, Edwards AN, Nawrocki KL, Anderson SE, Woods EC, McBride SM. The phosphotransfer protein CD1492 represses sporulation initiation in *Clostridium difficile*. Infect Immun. 2016;84(12):3434–44.

86. Sorg JA, Sonenshein AL. Bile salts and glycine as cogerminants for *Clostridium difficile* spores. J Bacteriol. 2008;190(7):2505–12.

87. Sorg JA, Sonenshein AL. Chenodeoxycholate is an inhibitor of *Clostridium difficile* spore germination. J Bacteriol. 2009;191(3):1115–7.

88. Francis MB, Allen CA, Sorg JA. Spore Cortex Hydrolysis Precedes Dipicolinic Acid Release during *Clostridium difficile* Spore Germination. J Bacteriol. 2015;197(14):2276–83.

